# Genome structure and population genomics of the canine heartworm *Dirofilaria immitis*

**DOI:** 10.1101/2023.04.25.538225

**Authors:** Javier Gandasegui, Rosemonde I. Power, Emily Curry, Daisy Ching-Wai Lau, Connor M. O’Neill, Adrian Wolstenholme, Roger Prichard, Jan Šlapeta, Stephen R. Doyle

**Affiliations:** Barcelona Institute for Global Health (ISGlobal), Hospital Clínic - University of Barcelona, Barcelona, Spain; Sydney School of Veterinary Science, Faculty of Science, The University of Sydney, NSW, Australia; Institute of Parasitology, McGill University, Sainte Anne-de-Bellevue, QC, Canada; Department of Infectious Diseases, University of Georgia, Athens GA 30602, USA; Wellcome Sanger Institute, Cambridgeshire, CB10 1SA, United Kingdom

**Keywords:** *Dirofilaria immitis*, whole-genome sequencing, genome assembly, population genomics, anthelmintic resistance

## Abstract

The heartworm, *Dirofilaria immitis*, is a filarial parasitic nematode responsible for significant morbidity and mortality in wild and domesticated canids. Resistance to macrocyclic lactone drug prevention represents a significant threat to parasite control and has prompted investigations to understand the genetic determinants of resistance. This study aimed to improve the genomic resources of *D. immitis* to enable a more precise understanding of how genetic variation is distributed within and between parasite populations worldwide, which will inform the likelihood and rate by which parasites, and in turn, resistant alleles, might spread. We have guided the scaffolding of a recently published genome assembly for *D. immitis* (ICBAS_JMDir_1.0) using the chromosomal-scale reference genomes of *Brugia malayi* and *Onchocerca volvulus*, resulting in an 89.5 Mb assembly composed of four autosomal- and one X-linked chromosomal-scale scaffolds representing 99.7% of the genome. Publicly available and new whole-genome sequencing data from 32 *D. immitis* samples from Australia, Italy and the USA were assessed using principal component analysis, nucleotide diversity (Pi) and absolute genetic divergence (Dxy) to characterise the global genetic structure and measure within- and between population diversity. These population genetic analyses revealed broad-scale genetic structure among globally diverse samples and differences in genetic diversity between populations; however, fine-scale subpopulation analysis was limited and biased by differences between sample types. Finally, we mapped SNPs previously associated with macrocyclic lactone resistance in the new genome assembly, revealing physical linkage of high-priority variants on chromosome 3, and determined their frequency in the studied populations. This new chromosomal assembly for *D. immitis* now allows for a more precise investigation of selection on genome-wide genetic variation and will enhance our understanding of parasite transmission and the spread of genetic variants responsible for resistance to treatment.

## 1. Introduction

The heartworm *Dirofilaria immitis* is a filarial nematode that primarily parasitises domestic and wild canids. Heartworm disease, or dirofilariasis, is caused by the presence of adult worms in pulmonary arteries that can cause inflammation, vascular dysfunction, and pulmonary hypertension, and circulating microfilariae in peripheral blood that cause acute and chronic inflammatory lesions in the lungs and other organs that, if not treated, can progress to congestive heart failure (Ames and Atkins, 2020). The mosquito-borne *D. immitis* can infect a wide range of other mammals and can be zoonotic (McCall et al., 2008), although human infections are relatively rare and are typically asymptomatic (Dantas-Torres and Otranto, 2013; Lee et al., 2010).

Prevention and control of dirofilariasis rely on preventive chemotherapy (PC) using anthelmintics, primarily macrocyclic lactones (ML) (McCall, 2005). Treatment of the definitive host aims to target the larval stages of the parasite after infection and prevents further development of the parasite to the pathogenic adult stages (Prichard, 2021). However, widespread and regular use of all anthelmintics places significant selection pressure on parasite populations and can promote the emergence of drug resistance (Kaplan and Vidyashankar, 2012; Rose Vineer et al., 2020). In the USA, reports describing the loss of efficacy of ML for the treatment of *D. immitis* infections have been increasing in frequency since 1998 (Hampshire, 2005) and confirmed by the isolation of an ML-resistant *D. immitis* isolate in 2014 (Pulaski et al., 2014). Several studies have attempted to define genetic variants associated with resistance to be used to monitor its emergence and spread (Bourguinat et al., 2017, 2016, 2011a, 2011b; Curry et al., 2022a; Mani et al., 2017; Sanchez et al., 2020; Willi et al., 2018). The most promising of these studies used pooled worm whole-genome sequencing of multiple phenotypically-defined sensitive and resistant isolates, which, after validation using independent genotyping, a set of 42 SNPs were proposed to be associated with poor treatment response and MLs resistance (Bourguinat et al., 2015). Subsequently, this initial set of genetic variants was re-assessed and reduced to 10- and two-SNP panels, validated using clinical samples, that were proposed to discriminate between susceptible and ML-resistant parasites (Ballesteros et al., 2018). This small panel of SNPs has been examined in European and Australian samples; however, additional support for these markers was not provided due to the lack of phenotypically-defined resistant isolates (Curry et al., 2022b; Power and Šlapeta, 2022). One caveat to the genome-to-SNPs approach was that the initial analyses were performed on a fragmented genome assembly and without a firm understanding of the genetic diversity within and between isolates, which can lead to both biological and technical biases that can confound association studies for genetic traits such as drug resistance (Doyle and Cotton, 2019). Improved genome resources and control of genetic diversity have been critical for increasing the genetic resolution to detect drug resistance loci in several helminths (Beesley et al., 2023; Doyle et al., 2022, 2019; Laing et al., 2022; Le Clec’h et al., 2021) and may aid not only in prioritising genetic markers but enable the identification of causal variants for ML resistance in *D. immitis*.

There are few studies focused on understanding the population genetics of *D. immitis*. The publication of the draft genome assembly nDi2.2 compared two isolates, one from Italy and one from the USA, revealing low levels of genetic divergence (Godel et al., 2012). Within the US, two studies based on microsatellite genotyping report variable levels of population structure and connectivity, from clustering individuals based on geographical origin (Belanger et al., 2011) to a more recent study that found a complete lack of genetic structure based on geographic origin or anthelmintic resistance phenotype (Sanchez et al., 2020). Although limited in number, these studies suggest that microsatellites are insufficient and that higher resolution data, such as whole genome sequencing, may be necessary to identify fine-scale genetic structure between *D. immitis* populations. A better understanding of population structure and connectivity of *D. immitis* in endemic regions will subsequently inform the likelihood and rate by which parasites, and in turn resistant alleles, might spread.

In this study, we have curated a recently published assembly to generate the first chromosomal assembly of *D. immitis*, providing new insight into autosome and sex-chromosome structure relative to closely related filarial nematodes. Using this new assembly, we have evaluated publicly available and new whole-genome sequencing data to understand the global population structure of *D. immitis*, from which we have defined levels of genetic diversity within and between populations. Finally, we have reassessed previously described genetic markers for ML resistance, now in a chromosomal context.

## 2. Materials and methods

### 2.1. Genome assembly

#### 2.1.1. Curating a chromosome-scale genome assembly

Our motivation to improve the genome assembly of *D. immitis* was driven by the need for high-quality contiguous genome resources to understand genome-wide genetic diversity, particularly in mapping genetic variation associated with drug resistance (Doyle et al., 2022; Doyle and Cotton, 2019). Until recently, the available genome of *D. immitis* was highly fragmented (nDi.2.2; n = 16,061 scaffolds)(Godel et al., 2012), limiting the interpretation of genome-wide variation to very discrete regions of the genome. The ICBAS_JMDir_1.0 assembly, generated from Nanopore long-read sequencing of DNA from a single worm (Gomes-de-Sá et al., 2022), significantly improved assembly contiguity (n = 110 contigs); however, the assembly was still fragmented, lacked chromosomal context and did not differentiate autosomal and sex-linked sequences. To improve the ICBAS_JMDir_1.0 assembly further, the genome was curated as follows.

First, we recognised that the ICBAS_JMDir_1.0 was missing a subset of universally conserved BUSCO genes (BUSCO v.5.4.3, nematoda ob10 lineage, n = 3,131 (Manni et al., 2021)) present in the draft nDi.2.2 assembly. In total, 26 contigs/scaffolds containing 30 complete BUSCO genes were recovered from nDi.2.2 and merged with the ICBAS_JMDir_1.0. Gaps in the newly added scaffold sequences were closed using PacBio reads (accession: SRR10533235) with TGS-GapCloser v.1.2.1 (Xu et al., 2020), reducing gaps from 468 to 275. The merged assembly was subsequently scaffolded using chromosomal assemblies of *Brugia malayi* (Foster et al., 2020) and *Onchocerca volvulus* (Cotton et al., 2016)(note: the post-publication “V4 assembly” of *O. volvulus* was used here, in which complete chromosomal scaffolds are resolved) as a reference using RagTag v2.1.0 (Alonge et al., 2022). This revealed greater sequence conservation between *D. immitis* and *O. volvulus* relative to *B. malayi*, consistent with the phylogenetic placement of these species (McCann et al., 2021). Two clear differences were observed between these intermediate assemblies: (i) that the chromosome number differed, with five chromosomal scaffolds after *B. malayi*-led scaffolding and four chromosomal scaffolds after *O. volvulus* led-scaffolding, and (ii) the structure of the sex-chromosome differed between *O. volvulus* and *B. malayi*, based on ancestral chromosome fusion and fission, with a conserved sex-specific region and divergent species-specific regions. Given that the *D. immitis* karyotype is 2n = 10, consistent with the chromosome number of *B. malayi*, we took the following approach: (i) the *O. volvulus*-led X chromosome (OVOC.OM2) was used to guide the delineation of the *D. immitis* X chromosome, and (ii) split the *O. volvulus* OM1a-led scaffold into two *D. immitis* chromosomes, Chr2 and Chr4, based on synteny with distinct *B. malayi* chromosomes. These decisions were supported by detecting intra-chromosomal telomere sequences within the intermediate but misassembled *B. malayi*-led X chromosome and *O. volvulus* OM1a-led scaffolds, which, when corrected, resulted in terminal telomere sequences in the curated *D. immitis* chromosomal sequences. The telomere location and frequency were determined using fastaq search_for_seq (https://github.com/sanger-pathogens/Fastaq) using a dimer of the telomeric monomer sequence “TTAGGC”. An additional round of gap-filling with TGS-GapCloser did not further reduce the number of gaps. Finally, redundant sequences were identified and removed with purge_dups v.1.2.5 (Guan et al., 2020).

#### 2.1.2. Wolbachia and mitochondrial genomes

*Wolbachia* contigs were removed from the *D. immitis* ICBAS_JMDir_1.0 assembly by the original authors (Gomes-de-Sá et al., 2022). To retrieve it for analysis here, raw Nanopore reads from the sample accession SRR14299255 were reanalysed. Porechop v.0.2.4 was first used to trim sequence adapters from the nanopore data before assembling the data with flye v.2.9.1-b1780 (parameters: --nano-raw --genome-size 100M). A single contig of 915,677 bp was assembled. We confirmed the *Wolbachia* genome by comparing it to the assembled genome from the Di2.2 assembly (wDi22; Dirofilaria_immitis_wolbachia_2.2.fna) using minimap2 v.2.16-r922. The mitochondrial genome was also missing from the ICBAS_JMDir_1.0. In this instance, the mitochondrial genome scaffold mDi_Athens_2.1 from the nDi.2.2 assembly was used.

#### 2.1.3. Genome comparison between Dirofilaria immitis assemblies and related nematodes

The assembly contiguity and completeness during assembly curation and relative to other *D. immitis* and related filarial nematodes was assessed in several ways. Genome assembly statistics were calculated using assembly-stats (https://github.com/sanger-pathogens/assembly-stats). Conserved BUSCOs were calculated using BUSCO v.5.4.3 (parameters: --mode genome --lineage_dataset nematoda_odb10; n = 3,131)(Manni et al., 2021). The *D. immitis* assemblies were compared by analysing the cumulative assembly size and sequence lengths generated using samtools faidx. Large-scale comparisons between genome assemblies were performed using minimap2 (parameter: -x asm20), from which paf outputs were visualised in R v.4.0.3 as a pairwise comparison of sequence hit positions and length (min: 300 bp) as dot plots or in a circular format showing links between chromosomes (min: 5,000 bp) using the R package circlize (https://github.com/jokergoo/circlize).

### 2.2. Samples

In total, data from 31 samples were used. Metadata regarding the samples are found in **Additional file 2: Table S1**.

#### 2.2.1. Publicly available datasets

A total of 12 publicly available datasets were used from the following ENA BioProject IDs, including (i) PRJNA681066: data from five adult samples from New South Wales (NSW), Australia (Lau et al., 2021); (ii) PRJNA588995: data from five microfilaria samples from the USA, including two from Georgia (GEO), one each from Illinois (ILL), Louisiana (LOU), and Mississippi (MIP) (Shin et al., 2020); and (iii) PRJEB2554: two adult samples, one from Georgia (GEO), USA, and one from Pavia (PAV), Italy (Godel et al., 2012).

#### 2.2.2. Additional, unpublished datasets

An additional 19 datasets were curated from collaborators, including:

(i) twelve pooled microfilaria samples from Queensland (QLD), Australia. These Australian blood samples (32 to 38,250 microfilaria.ml^-1^) and their DNA were included in a previous study of ML resistance in Australia (Power and Šlapeta, 2022). Extracted DNA from filtered microfilariae of *D. immitis* from 2019 (HW3/19, HW4/19) and 2020 (HW5/20 to HW9/20, HW12/20 to HW16/20) were sent to Novogene (HK) Co., Ltd for indexing, library construction and whole-genome sequencing; Illumina NovaSeq sequencing was performed at an expected depth of 6G with 150 bp paired-end reads.

(ii) Seven pooled microfilaria samples from the USA, including two from Georgia, and one each from Arkansas (ARK), Louisiana (LOU), Michigan (MCH), Tennessee (TEN), and Texas (TEX). DNA libraries were sequenced on a HiSeq X10 using 150 bp paired-end reads.

### 2.3. Genomic analyses: data preparation, mapping & variant calling

#### 2.3.1. Evaluation of sample contamination

Given the variable sample provenance and heterogeneity, the degree of contamination in the raw sequencing reads was evaluated using Kraken2 v.2.1.2 (Wood and Salzberg, 2014). The raw reads were analysed using two approaches; (i) using the minikraken2 8Gb database to assess sample contamination with bacteria and/or viruses, and (ii) using a custom Kraken database consisting of the *D. immitis* (nDi.2.2) and *the* domestic dog *Canis lupus familiaris* (GenBank accession: GCA_014441545) genomes. The output of Kraken2 was summarised using MultiQC v.1.14 (Ewels et al., 2016).

#### 2.3.2. Trimming, mapping and processing of sequencing data

Quality-based read trimming was performed to remove poor-quality reads and contaminating adapter sequences using trimmomatic v.0.39 (Bolger et al., 2014) (parameters: ILLUMINACLIP:Illumina-adapters.fa:2:30:10 SLIDINGWINDOW:10:20 MINLEN:50). Trimmed reads were mapped to the improved *D. immitis* reference genome (supplemented with mitochondrial and *Wolbachia* genomes) using minimap2 v.2.16-r922 (Bolger et al., 2014; Li, 2018). Sambamba v.0.6.6 was used to mark duplicate reads. For samples with multiple read sets, mapped reads were combined using samtools merge (v.1.14).

#### 2.3.3. Genome coverage analysis

Sequencing coverage throughout the genome was determined using a combination of bamtools (Barnett et al., 2011), bedtools makewindows (v2.29.0) (Quinlan, 2014), and samtools bedcov (v.1.14)(Li et al., 2009), in sliding windows and per chromosome. The ratio of X chromosome to autosome coverage was used to infer sample sex. Coverage data of nuclear, mitochondrial and *Wolbachia* genome coverage are shown in **Additional file 2: Table S2**. Samples from Queensland showed very low nuclear and *Wolbachia* coverage, whereas mitochondrial coverage was higher in comparison. This can be expected since the number of mitochondria per cell increases the number of DNA copies, which facilitates mapping its genome. However, in all cases, coverage was highly variable since sample collection, sample type, and sequencing approaches differed between samples.

#### 2.3.4. Variant calling and filtering

Variant calling was performed using GATK v.4.1.4.1 (Van der Auwera and O’Connor, 2020). Initially, variants were identified, and GVCF files were generated for each BAM file using the tool *HaplotypeCaller*. Then variants were consolidated, and GVCF files merged using *CombineGVCF*s before joint-call cohort genotyping using GATK *GenotypeGVCFs*. Finally, a single multisample VCF file containing all initial variants and samples was generated.

Variant filtering was performed using GATK *SelectVariants* and *VariantFiltration*. Nuclear, mitochondrial and *Wolbachia* variants were filtered separately and following the same workflow. A data-led approach was employed to determine the distribution of quality metrics, including QUAL, DP, MQ, SOR, FS, QD, MQRankSum, and ReadPosRankSum, by removing relevant tails of variant distributions (typically the upper and/or lower 1-5%).

Variants were further filtered using vcftools v.0.1.16 (Danecek et al., 2011) to ensure they met the following criteria: minimum and maximum alleles = 2; minor allele frequency (MAF) > 0.02; Hardy Weinberg Equilibrium = >1E-6 (nuclear variants) and per genotype depth > 3. Considering the MAF threshold and our sample size, only variants with at least two alleles in the population were considered. Per-sample missingness was evaluated to identify samples with high levels of missing data. Finally, per-site missingness was evaluated to establish a common threshold for all subsets of variants. Only SNPs located in chromosome 1 to chromosome 4 were retained for the population genetic analyses, excluding variants in the shorter scaffolds (which constitute ∼0.3% of the genome) and the sex-linked X chromosome.

Due to per-sample missingness, all samples from QLD had to be removed from the nuclear variant set (**Additional file 1: Fig. S4a)**, and two samples from QLD were removed from the mitochondrial variant sets (*Additional file 1: Fig. S4b*), whereas all samples from QLD and three from the USA (ARK, TEN and TEX) were removed from the *Wolbachia* subset of variants ***(Additional file 1: Fig. S4c)***. Therefore, 19 samples were assessed using nuclear variants, 29 samples with mitochondrial variants, and 18 with *Wolbachia* variants. A per-site missingness threshold of 0.8 for nuclear and mitochondrial sets and 0.7 for the *Wolbachia* variant subset was applied (see code repository for more information about the filtering process).

### 2.4. Genomic analyses: population genomics

#### 2.4.1. Principal component analysis

Given the mixture of pooled and single worm sample types, broad-scale genetic relatedness between samples and populations was explored using allele frequencies rather than genotypes. Allele frequencies were extracted using the R package vcfR (Knaus and Grünwald, 2017) from the AD (allele depth) parameter in the multisample VCF file as:

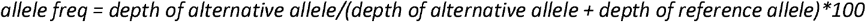

PCA on allele frequencies was performed using the R function prcomp(). It is important to note that prcomp() does not allow missing values; thus, 126,728 nuclear SNPs were included in this analysis. PCA of mitochondrial and *Wolbachia* variants was performed using the R package SNPRelate (Zheng et al., 2012) instead of allele frequencies, which allowed us to incorporate missing SNPs in the PCA and thereby use more variants in the analysis.

To explore the clustering of USA samples in the PCA of nuclear variants, other parameters and technical aspects potentially driving the observed clustering of USA samples were assessed. The depth per sample was obtained using the R package vcfR (Knaus and Grünwald, 2017), whereas the missingness and the excess of heterozygosity (inbreeding coefficient “F”) were estimated using vcftools on nuclear variants.

#### 2.4.2. Genetic diversity within and between populations

Genome-wide nucleotide diversity (Pi) within populations and absolute nucleotide divergence (Dxy) between pairs of populations were determined in 100 kb non-overlapping sliding windows using pixy v.1.2.7.beta1 (Korunes and Samuk, 2021). Pixy is a command line tool for calculating the genetic diversity summary statistics that account for invariant sites, which are essential for the correct computation of Pi and Dxy. Briefly, all individual GVCF files were genotyped using the tool GenotypeGVCFs with the --all-sites flag, which allows the inclusion of invariant sites during genotyping. First, invariant sites were selected using the flag --select-type-to-include ‘NO_VARIATION’ in the GATK tool SelectVariants and then merged with the previously filtered nuclear SNPs set (see section *Variant calling and filtering*). As mentioned above, only nuclear positions in the main chromosomes were selected. SNPs and invariants were merged using the GATK MergeVcfs function and indexed using samtools.

#### 2.4.3. Mapping of proposed genetic markers for macrocyclic lactone resistance

Markers for anthelmintic resistance were originally developed using the fragmented nDi.2.2 genome. They were distributed on many different contigs and scaffolds with little understanding of how they related to each other in the genome. To address this, SNP coordinates and 100 bp flanking region were extracted from the nDi.2.2 using bedtools get_fasta and mapped to the dimmitis_WSI_2.2 genome using nucmer v.3.1 (Kurtz et al., 2004). Of the 42 original markers, two markers–NODE_42411 (nDi.2.2.scaf00001:466197) and NODE_9400 (nDi.2.2.scaf00005:662854)–were not mapped. The above mapping process was repeated, but this time, using a 10,000 bp flanking region to increase the likelihood of finding the region of the genome; however, the sites still could not be found, leading us to conclude that they were redundant sequences in the original assembly that were removed during curation. SNP positions in the curated genome were confirmed by identifying variants in the variant call format (VCF) file from the population genomics data, from which allele frequencies per population were calculated.

## 3. Results and Discussion

### 3.1. Genome structure of Dirofilaria immitis

Comparative karyotype studies from male and female parasites indicate the *D. immitis* genome is composed of four autosomes, with heterogametic males containing additional X and Y sex chromosomes and homogametic females containing two X chromosomes (2n=10) (Delves et al., 1986; Sakaguchi et al., 1980). This karyotype is broadly consistent with the related filarial nematodes *Brugia malayi, Wuchereria bancrofti*, and some *Onchocerca* spp. (excluding *O. volvulus* and *O. gibsoni*, which have undergone an additional chromosomal fusion to produce 3A + XY) (Post, 2005).

Recently, a high-quality *de novo* assembly (ICBAS_JMDir_1.0) for *D. immitis* was generated using long-read Oxford Nanopore sequencing of DNA derived from a single parasite (Gomes-de-Sá et al., 2022), representing a significant improvement in genome contiguity relative to the original draft assembly, nDi2.2, published in 2012 (Godel et al., 2012) ***(Table 1)***. To provide greater insight into the genome structure of *D. immitis* and to better understand genome-wide genetic variation among globally diverse isolates, we guided scaffolding of the ICBAS_JMDir_1.0 assembly against the chromosomal reference genomes of *B. malayi* and *O. volvulus* **(Fig. 1a and Fig. 1b**, respectively). This approach achieved an assembly containing 99.7% of sequence data in four autosomal scaffolds of approximately equal length and one X-linked chromosomal-scale scaffold approximately twice the length of the autosomes, consistent with the expected female karyotype and distribution of conserved Nigon units of chromosome structure (Gonzalez de la Rosa et al., 2021). Although not all contigs were incorporated into chromosomal scaffolds and gaps have been introduced by scaffolding, relative to previous assemblies, this curated assembly is highly contiguous (**Additional file 1: Fig. S1)** and is more complete based on a comparison of conserved BUSCO sequences (**Table 1)**. The X chromosome was identified based on synteny with the *O. volvulus* X chromosome and half coverage on the *D. immitis* X chromosome from sequencing of male parasites (**Fig. 1c)**. The presence of both half and full coverage on the X chromosome is consistent with a shared pseudoautosomal region of the X and Y chromosomes, with the half-coverage region representing the X-chromosome-specific region; we could not resolve the Y-chromosome-specific region, which may be achieved by further long-read sequencing of male parasites. Defining the sex-specific region on X enabled the delineation of male (X-to-autosome [X:A] ratio ≈ 0.5) and female samples (X:A ≈ 1.0) from mixed-sex populations, i.e., pooled microfilaria and adult females with gametes (X:A ≈ 0.75), based on sequencing data alone.

**Table 1.** Genome assemblies of *Dirofilaria immitis* and comparators with other filarial genome assemblies.

|  | *D. immitis* | *D. immitis* | *D. immitis* | *Brugia malayi* | *Onchocerca volvulus* |
| --- | --- | --- | --- | --- | --- |
| Version | dimmitis_WSI_2.2; this study | ICBAS_JMDir_1.0; (Gomes-de-Sá et al., 2022) | nDi.2.2; (Godel et al., 2012), WBP17 | WBP17; (Foster et al., 2020) | V4; WBP17 (Cotton et al., 2016) |
| Assembly size (bp) | 89,475,811 | 86,813,779 | 88,309,529 | 88,235,797 | 96,427,137 |
| Number of scaffolds (n) | 27 | 110 | 16,061 | 197 | 708 |
| Average length (bp) | 3,313,918.93 | 789,216.17 | 5,498.38 | 447,897.45 | 136,196.52 |
| Largest scaffold (bp) | 28,232,375 | 12,535,412 | 1,085,577 | 24,943,668 | 28,345,163 |
| Scaffold N50 (bp) | 15,199,470 | 4,253,153 | 71,281 | 14,214,749 | 25,485,961 |
| Scaffold N50 (n) | 3 | 7 | 219 | 3 | 2 |
| Number of gaps (n) | 156 | 0 | 13,431 | 8 | 581 |
| Number of Ns (bp) | 40,261 | 0 | 3,430,609 | 277,365 | 3,074,367 |
| BUSCO completeness<sup>1</sup> | C:94.3 [S:93.0, D:1.3], F:1.7, M:4.0 | C:94.1 [S:93.7, D:0.4], F:1.6, M:4.3 | C:93.7 [S:91.5, D:2.2], F:1.8, M:4.5 | C:98.5 [S:97.9, D:0.6], F:0.6, M:0.9 | C:97.9 [S:97.3, D:0.6], F:1.0, M:1.1 |

1. BUSCO v.5.4.3 was run using nematoda\_odb10 lineage (n = 3,131); percentage of total BUSCOs shown; C = complete, S = complete, single-copy, D = complete, duplicated, F = fragmented, M = missing.

**Fig. 1.**
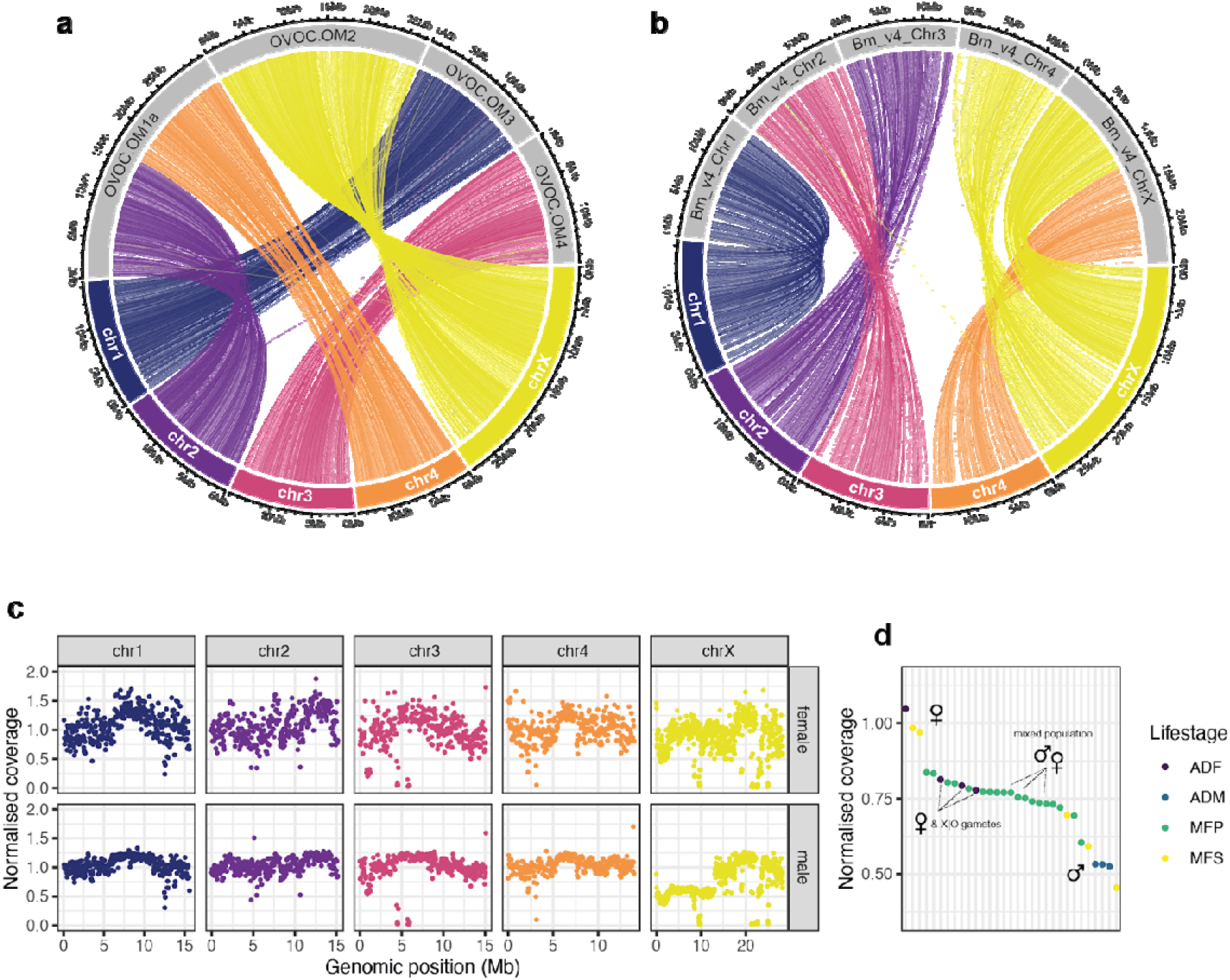
Genome structure and coverage of Dirofilaria immitis chromosomes. Broad-scale chromosome structure and rearrangements between the curated *D. immitis* genome dimmitis_WSI_2.2 and (*a*) *Onchocerca volvulus* and (*b*) *Brugia malayi* chromosomal assemblies is shown. Each link between chromosomes represents a minimum 5 kb hit determined by whole genome alignments using minimap2. (*c*) Normalised sequencing coverage of the *D. immitis* chromosomes comparing a female (top) and male (bottom) sample to highlight the drop in coverage in the first half of the X chromosome in the male sample. Each point represents coverage per 50 kb window throughout the genome, each normalised by the genome-wide median coverage. (*d*) Comparison of normalised X-to-autosome (X:A) coverage among the sample cohort. ADF = adult female, ADM = adult male, MFP = pooled microfilaria, MFS = single microfilaria.

Although we identified regions of conserved synteny with both *B. malayi* and *O. volvulus* genomes, considerable rearrangements, particularly toward the ends of the chromosomes, were also identified **(Additional file 1: Fig. S2)**. We note that without independent evidence derived from *D. immitis* to order and orientate the assembly, these scaffolds should be treated cautiously and may contain assembly errors from the original nanopore assembly and subsequent reference-based scaffolding. Nonetheless, these chromosome-scale scaffolds represent conserved linkage groups based on synteny against both *O. volvulus* and *B. malayi* chromosomal assemblies, delineating autosome and sex chromosomes and providing a means to differentiate chromosome-specific genetic markers.

### 3.2. Genome-wide genetic variation within and between broadly sampled populations

The genome-wide distribution of genetic variation from 32 *D. immitis* datasets was assessed (see **Additional file 2: Table S1** for details). The samples from which these data were derived were heterogeneous regarding biological stage and sample type: seven came from single adult worms, five from single microfilaria and 19 from pooled microfilaria. Considering the limited sample availability, the geographic range of sampling was also limited to three regions **(Fig. 2a):** 17 samples were collected in Australia (12 from QLD and five from NSW), one sample was collected in Pavia (PAV), Italy, and the remaining 13 samples were collected in the USA (one sample in Michigan, Illinois, Tennessee, Arkansas, Mississippi and Texas; five samples in Georgia and two samples in Louisiana).

**Fig. 2.**
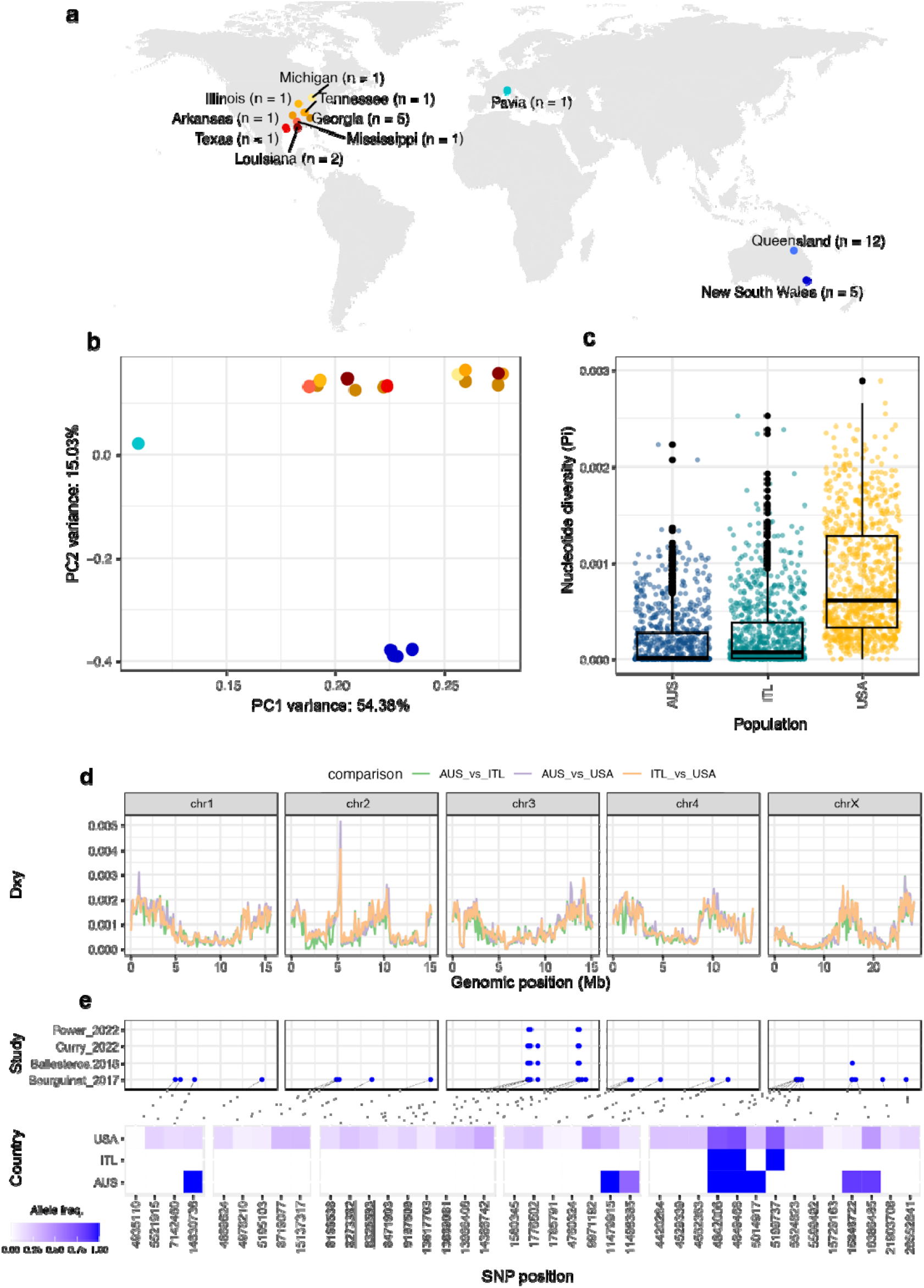
Population structure and genome-wide analyses of within and between population genetic diversity. (**a**) A world map shows the broad locations (city/state) from which samples analysed were originally collected. Sample numbers per location are indicated. (***b***) Principal component analysis (PCA) using 126,728 nuclear SNP frequencies from 19 samples (Additional file 1 explains the details about the variant filtering process). PC1 and PC2 are shown, which explain 54.4% and 15.0% of the variation, respectively. Samples are coloured by location shown in panel (a). **(*c*)** Box plots show nucleotide diversity (Pi) distribution per geographic region. Each point represents 100 kb non-overlapping sliding windows in the genome. (***d*)** Comparison of absolute nucleotide divergence (Dxy) genome-wide between pairs of populations. Colours indicate paired comparisons, including AUS vs ITL (green), AUS vs USA (purple), and ITL vs USA (orange). Vertical dashed lines indicate chromosome boundaries. *(e)* Chromosomal placement of genetic markers previously proposed to be associated with macrocyclic lactone resistance (top panel). The original 42 markers were developed using the fragmented assembly nDi.2.2 (Bourguinat_2017); here, we show some of these variants cluster on discrete regions of chromosomes. Subsequent studies that refine (Ballesteros_2018; 10 and two SNP models) and validate these SNPs in independent populations (Curry_2022 and Power_2022) are shown. The allele frequencies of the variants in the study populations are shown (bottom panel), with the 10 SNP model variants in bold and the two SNP model varisants in bold and underlined. Abbreviations: AUS = Australia; ITL= Italy; USA = United States of America.

Genetic diversity was assessed in the three genomes of *D. immitis*: the parasite’s nuclear and mitochondrial genomes and its intracellular endosymbiont, *Wolbachia*. In total, we identified 422,935 nuclear single nucleotide variants (of which 96,313 were indels), 49 mitochondrial variants (incl. ten indels) and 447 *Wolbachia* variants (incl. 198 indels). Due to the variable amount of sequencing data, mapped read coverage and levels of contamination (i.e. dog and bacterial DNA; **Additional file 1: Fig. S3**) per sample, we subset groups of samples for each analysis. Analyses of broad-scale genetic relationships between samples using principal component analysis (PCA) revealed only weak genetic structure among samples using either mitochondrial or *Wolbachia* variants alone **(Additional file 1: Fig. *S5a*** and **Fig. S5b**, respectively). The lack of genetic structure was likely due to the low number of mitochondrial variants and the high and variable levels of missing data in *Wolbachia* variants **(Additional file 1: Fig. S4c)**. Conversely, PCA using nuclear variants showed a broad-scale population structure clearly separating samples belonging to the three sampling countries (Australia, Italy and the USA) **(Fig. 2b)** with ∼70% of the genetic variance explained by the first two principal components. Furthermore, two distinct clusters of samples were observed among the samples from the USA. However, the two clusters of samples were not divided by sampling location or other geographic variables, as might be expected.

To understand these two USA clusters and differentiate potential technical from biological factors that might be driving this signal in the PCA, we assessed additional variables, including the sample type (pooled vs single worm), sequencing depth, per sample missingness, and heterozygosity estimated by inbreeding coefficient F in the PCA. These analyses highlighted that sample type–whether the sample was obtained from a single worm or a pooled population of worms, i.e. pooled microfilaria–more obviously explained the two clusters than geography. Only a single pooled sample (TEX, strong red colour in **Additional file 1: Fig S6a)** is grouped with data from single individuals. This sample had lower coverage when compared to the other pooled samples **(Additional file 1: Table S2)**, which may have consequently resulted in its grouping with other low-coverage, single-worm samples. The pooled-versus-single worm effect is correlated with sequencing depth (higher depth in pooled samples; **Additional file 1: Fig. S6b)**, missingness (less missing data in pooled samples; **Additional file 1: Fig. S6c)**, and heterozygosity (excess heterozygotes in pooled samples and excess homozygotes in single worm samples; **Additional file 1: Fig. S6d)**. These data highlight the challenge of working with mixed sample types and that differences in sample type (single/pooled, microfilaria/adults) and sample processing (DNA extraction, sequencing platform/depth) will impact the ability to define population genetic structure and genetic associations precisely. Such biases are important when comparing genetic traits like anthelmintic resistance when sampling has been performed across different hierarchical levels (e.g., life stages, within/between countries) (Doyle and Cotton, 2019).

To explore within and between population variation, we calculated nucleotide diversity (Pi) and nucleotide divergence between populations (Dxy) in 100 kb non-overlapped sliding windows throughout the genome. Genome-wide, Pi tended to increase towards the ends of chromosomes **(Additional file 1: Fig. S7)**, which is best demonstrated on chromosomes 1, 3 and X, the latter of which has an intrachromosomal peak in Pi that likely reflects the ancient chromosome fusion shared with *O. volvulus* (Cotton et al., 2016). Pi was significantly different between populations (p<0.001; Wilcoxon rank sum test), being highest in samples from the USA (mean[Pi] = 7.36e-4), followed by the Italian sample (mean[Pi] = 8.6e-5) and then the Australian samples (mean[Pi] = 3.9e-5) **(Fig. 2c)**. These values of genetic diversity are relatively low when compared to other vector-borne filarial parasites, such as *O. volvulus* (mean[Pi] = 4.0e−3) (Choi et al., 2016) and *W. bancrofti* (mean[Pi] = 2.4e-4) (Small et al., 2016). This low level of nucleotide diversity may limit the rate and likelihood of resistance to treatment developing broadly, or at least relative to some livestock-infective helminths in which resistance is widespread and evolved rapidly (Samson-Himmelstjerna et al., 2021), likely as a consequence of high levels of genetic diversity (Doyle et al., 2020). We also explored diversity within the USA populations, first considering the state where the sample was collected and then the sample type (pooled or single specimens). These data suggest that some populations are more diverse than others in the USA **(Additional file 1: Fig. S8a);** however, this interpretation is somewhat confounded by the fact that there are different sample types per population and that pooled samples showed higher diversity than single samples **(Additional file 1: Fig. S8b)**. Therefore, these data further support the relevance of the preparation of the samples before sequencing in the assessment of population structure and genetic epidemiology. When more sequencing data becomes available not only for *D. immitis* but also for other helminth parasites, future work must evaluate different approaches to integrate large datasets of specimens with different origins and life stages. Finally, the lack of apparent genetic structure and higher nucleotide diversity among geographically defined samples within the USA suggests a largely panmictic population and widespread transmission zone despite being a vector-transmitted disease, which, if true, entails significant challenges to ongoing control. Divergence in nucleotide diversity, Dxy, between populations showed similar patterns to genome-wide Pi throughout the chromosomes with increased variance toward the ends of chromosomes and small discrete peaks of differentiation throughout **(Fig. 2d)**. There was, however, a subtle trend to suggest AUS_vs_ITL was less divergent and AUS_vs_USA more divergent than USA_vs_ITL.

### 3.3. Chromosomal context of genetic markers associated with ivermectin resistance

Resistance to macrocyclic lactones is increasingly recognised as a threat to *D. immitis* control, particularly in the USA, where drug failures and resistant isolates have been defined and characterised (Diakou and Prichard, 2021; Maclean et al., 2017; Pulaski et al., 2014). Understanding the genetic determinants of drug resistance has been challenging, partly due to a lack of high-quality genome resources and a poor understanding of how genetic variation is distributed within and between populations under drug pressure. A panel of 42 candidate resistance genetic markers derived from whole genome sequencing data were identified (Bourguinat et al., 2015), which was subsequently refined to a small panel (between two and ten SNPs) that were predictive of a resistance phenotype (Ballesteros et al., 2018; Bourguinat et al., 2017; Curry et al., 2022b). However, these genetic markers were identified and developed using the fragmented draft genome nDi.2.2, and, therefore, their positions in the genome relative to each other–which may provide insight into the mode of drug selection and degree of linked selection–were largely undefined.

We have mapped 40 of 42 SNP positions on the chromosomes, revealing hits to all five chromosomes (**Fig. 2e;** top panel). Surprisingly, some variants were found in discrete clusters, including those proposed to be most predictive of resistance; notably, on chromosome 3, six SNPs from three distinct nDi.2.2 scaffolds were found in one cluster (range: 8166538-9197509 bp), and in another cluster, four SNPs from three distinct nDi.2.2 scaffolds (range: 13617703-14388742 bp) were mapped **(Fig. 2e)** These two clusters on chromosome 3 contained all but one mapped SNP from the two and ten SNP predictive models defined by Ballesteros and colleagues (Ballesteros et al., 2018). Therefore, the chromosomal context of the new genome has provided insight into the physical linkage between these specific variants, which could not be achieved using the draft genome alone. Analysis of allele frequency in our studied populations showed that these variants are present at a moderate frequency in the USA but are largely absent or are fixed variants unlikely to be associated with resistance from Italy and Australian samples (**Fig. 2e;** bottom panel), consistent with previous studies (Curry et al., 2022b; Power and Šlapeta, 2022). The relevance of these predictive variants outside of the USA is unknown and requires further validation with phenotypically defined non-USA isolates. We are underpowered to further resolve the genetics of resistance based on low sample numbers and limited drug susceptibility data among the available samples analysed here. However, these results demonstrate the value of a curated chromosomal genome assembly (Doyle, 2022), enabling a more precise differentiation between genetic variants associated with drug selection from unlinked background variation. Furthermore, it provides a scaffold for genetic mapping and genome-wide association experiments in *D. immitis*, which has been essential for understanding the genetics of anthelmintic resistance in other helminth species of humans (Doyle et al., 2017; Le Clec’h et al., 2021) and veterinary importance (Beesley et al., 2023; Doyle et al., 2022).

## 4. Conclusion

We have curated a contiguous and largely complete genome assembly for *D. immitis*, revealing the chromosome structure and genomic rearrangements for the first time. Further validation of assembly joins and structural variation using orthogonal sequencing technologies, for example, HiC chromatin conformation capture, will ensure this chromosomal assembly remains a valuable resource for the parasitology community for some time. Using publicly available genome data supplemented with new sequence data, we have described broad-scale within- and between-population genetic diversity among geographically diverse isolates. Available data limit our resolution toward understanding genome-wide diversity and differentiation between populations but will improve by denser sampling and sequencing of parasites throughout their natural range. Finally, a chromosomal assembly now allows a more precise investigation of selection on genome-wide genetic variation and will enhance our understanding of parasite transmission and the spread of genetic variants responsible for resistance to treatment.

## Abbreviations

AD: allele depth
BUSCO: Benchmarking Universal Single-Copy Orthologs
Dxy: absolute nucleotide divergence
PC: preventive chemotherapy
PCA: principal component analysis
MAF: minor allele frequency
ML: macrocyclic lactone
Pi: nucleotide diversity
SNP: single nucleotide polymorphism
VCF: variant call format
X:A: X-to-autosome ratio

## Declarations

### Ethics approval and consent to participate

Dog sampling in Australia was obtained after receiving written consent from the owners, and research performed at the University of Sydney was approved by the Animal Research Ethics Committee, which follows the NSW Animal Research Act 1985 and the Australian code for the care and use of animals for scientific purposes 8th edition (2013).

### Consent for publication

Not applicable

### Availability of data and materials

Raw sequencing data used in this study are publicly available from the European Nucleotide Archive (ENA) and are described in Table S1. The code used to generate and analyse data and plot figures can be found at https://github.com/Gandasegui/pop_genomics_dirofiliaria. The dimmitis_WSI_2.2 genome is available at: ftp://ngs.sanger.ac.uk/production/pathogens/sd21/dimmitis_genome/dimmitis_WSI_2.2.fa

### Competing interests

The authors declare that they have no competing interests.

### Funding

ISGlobal receives support from the Spanish Ministry of Science and Innovation through the “Centro de Excelencia Severo Ochoa 2019-2023” Program (CEX2018-000806-S) and support from the Generalitat de Catalunya through the CERCA Program. SRD is supported by a UKRI Future Leaders Fellowship [MR/T020733/1] and the Wellcome Trust through core funding to the Wellcome Sanger Institute [206194]. The Canine Research Foundation and Dogs Victoria, Australia, supported the research in Australia. For the purpose of Open Access, the author has applied a CC BY public copyright licence to any Author Accepted Manuscript version arising from this submission.

### Authors’ contributions

SRD designed and conceptualised the study. RIP, DC-WL, EC, CMO, AW, RP, and JŠ provided samples, sequencing data and parasitology expertise. JG and SRD carried out bioinformatic analyses. JG and SRD wrote the original draft manuscript, which was subsequently revised by all authors. All authors read and approved the final manuscript.

## Acknowledgements

We gratefully acknowledge the Pathogen Informatics group at the Wellcome Sanger Institute for informatics support, the Doyle group for constructive feedback on the analyses and manuscript, and the parasitology community for their open access data and sharing of genome resources.

## Supplementary information

**Additional file 1**: Supplementary Figures. **Fig S1**. Comparison of genome assemblies of *Dirofilaria immitis*. **Fig S2**. Chromosomal synteny between *Dirofilaria immitis* and related filarial species. **Fig S3**. Contamination and percentage of *Dirofilaria immitis* and *Canus lupus* DNA estimated by Kraken. **Fig S4**. Per-sample missingness for each subset of variants. **Fig S5**. PCA on mitochondrial (a) and *Wolbachia* (b) variants. **Fig S6**. PCA of nuclear variants from USA samples. **Fig S7**. Genome-wide patterns of nucleotide diversity between USA, Italian and Australian samples. **Fig S8**. Nucleotide diversity (Pi) within the USA.

**Additional file 2: Table S1**. Sample origin and geographical information. **Table S2**. Genome mapping data by sample, including nuclear, mitochondrial and *Wolbachia* genome coverage. **Table S3**. Mapping resistance-associated variants to the chromosomal genome assembly.

## Notes

### Competing Interest Statement

The authors have declared no competing interest.

