## Additional file 1 for "Genome structure and population genomics of the canine heartworm *Dirofilaria immitis*"

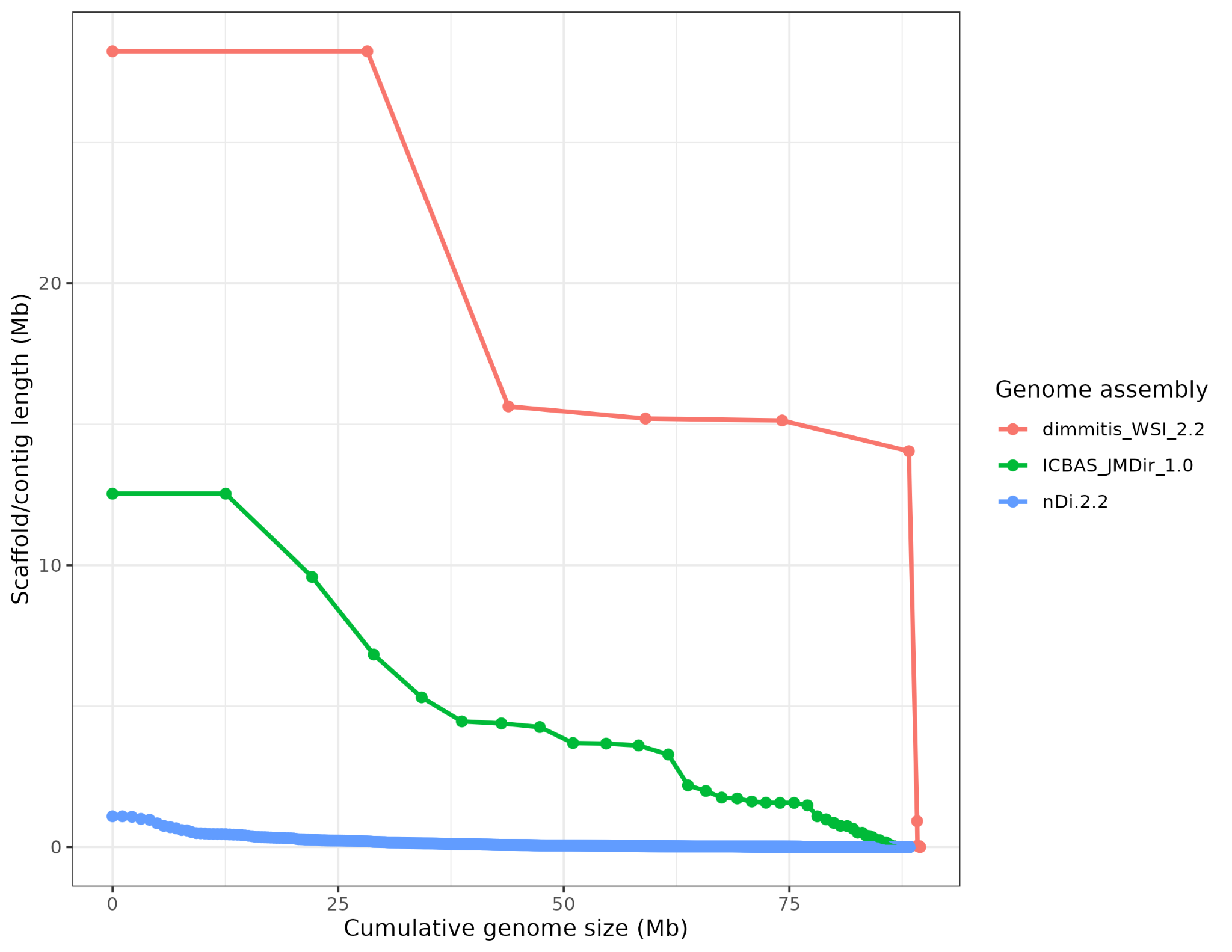

**Fig. S1. Comparison of genome assemblies of *Dirofilaria immitis*.** The plot shows the relationship between scaffold/contig length and cumulative genome size between the original draft assembly nDi.2.2, the greatly improved Nanopore assembly ICBAS_JMDir_1.0, and the curated assembly described in this study, dimmitis_WSI_2.2. Each scaffold/contig in the assemblies is represented as a single point, sorted by length. Although the overall genome size is relatively consistent between assemblies (86-89 Mb), the number and length of sequences are markedly different.

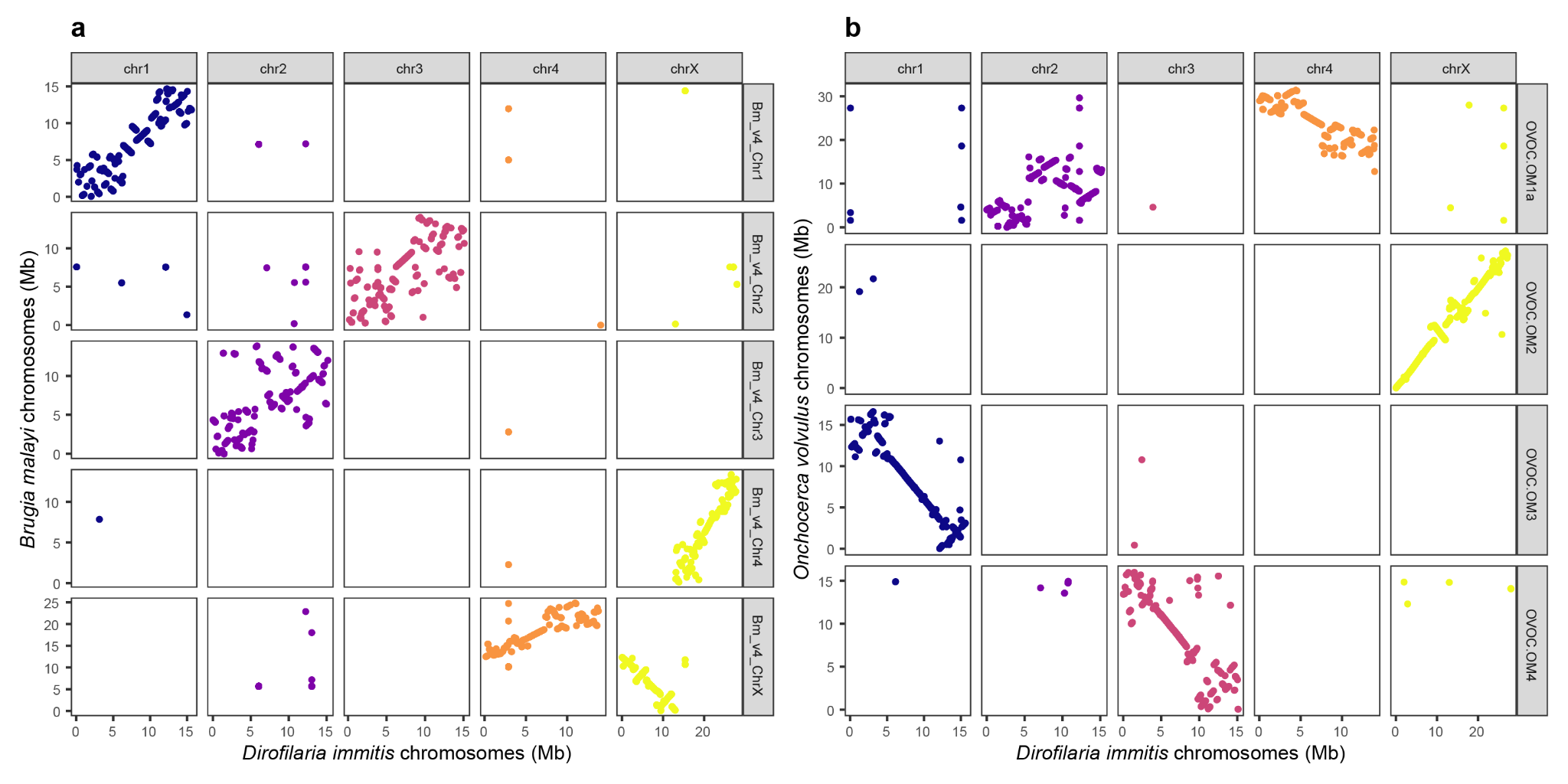

**Fig. S2. Chromosomal synteny between *Dirofilaria immitis* and related filarial species.** Dot plots show the relationship between the curated

*D. immitis* genome and the chromosomal genome assemblies of (**a**) *Brugia malayi* and (**b**) *Onchocerca volvulus*. In both plots, each point represents a minimum 5 kb sequence match between pairs of genomes determined using minimap2. The degree of synteny is indicated by the length of the adjacent points forming a line indicating the degree of synteny. These plots highlight that most of the chromosomal content is conserved between ancestral chromosomes and that *D. immitis* has greater synteny with *O. volvulus* than *B. malayi*, with the exception that *D. immitis* X is comprised of Chr4 and ChrX of *B. malayi*, and *D. immitis* Chr2 and Chr4 are joined to form OVOC.OM1a of *O. volvulus*. We note that some of these arrangements will be driven by misassemblies and need validation using an independent sequencing approach such as HiC chromatin conformation capture.

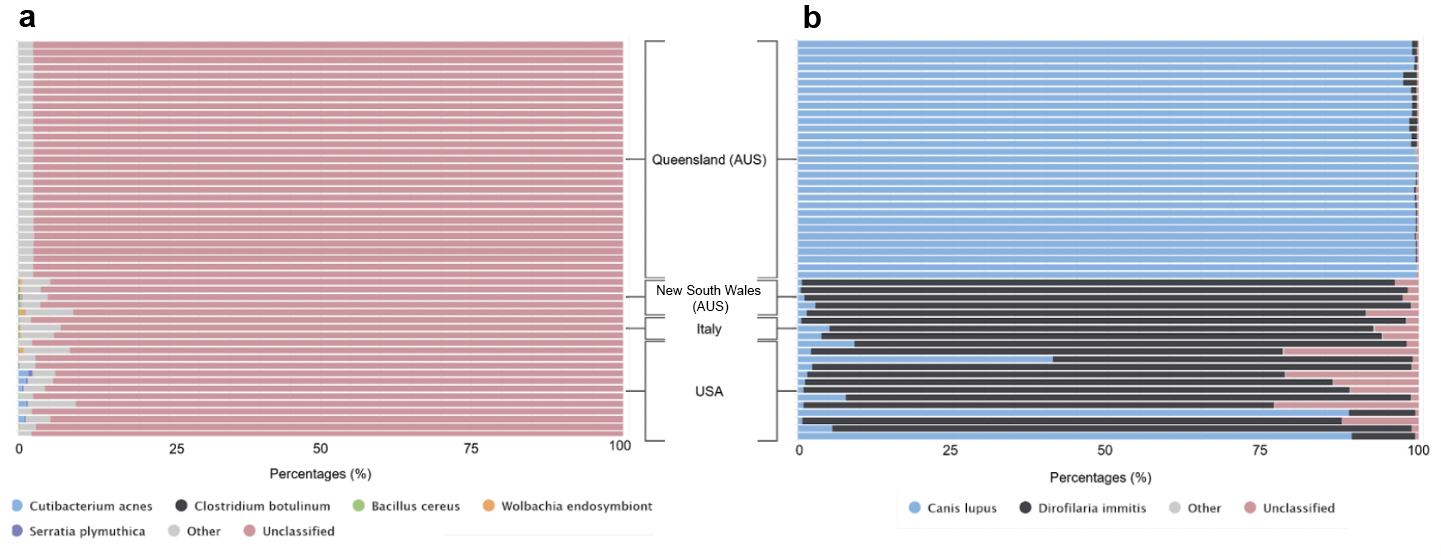

**Fig. S3. Contamination and percentage of *Dirofilaria immitis* and *Canus lupus* DNA estimated by Kraken.** (**a**) Percentage sample contamination with bacteria and/or viruses using minikkraken 8Gb database. (**b**) Percentage of *D. immitis* and *C. lupus* DNA reads. Overall, the degree of contamination from bacteria and/or viruses was low in all samples; however, the proportion of *D. immiti*s and *C. lupu*s DNA reads differed significantly between sample datasets. All Queensland samples had low counts of *D. immitis* sequencing reads and a high proportion of reads derived from the dog genome, which is consistent with the fact that these samples were pooled microfilaria isolated from dog blood.

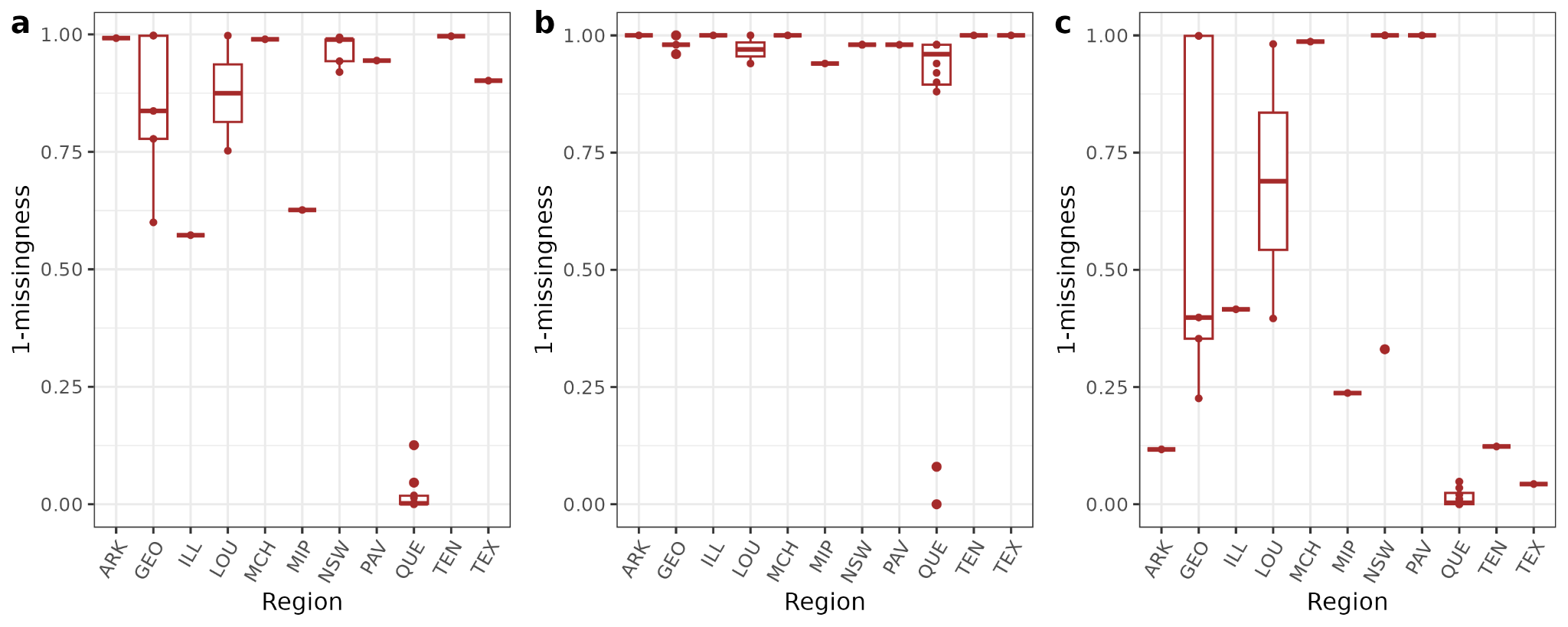

**Fig. S4.** **Per-sample missingness for each subset of variants.** (**a)** Nuclear, (**b**) mitochondria, and (**c**) Wolbachia variants. The y-axis represents the proportion of variants estimated by 1-missingness, whereas the x-axis shows the populations at the regional level. Abbreviations: ARK, Arkansas; GEO, Georgia; ILL, Illinois; LOU, Louisiana; MCH, Michigan; MIP, Mississippi; NSW; New South Wales; PAV, Pavia; QUE, Queensland; TEN, Tennessee; TEX, Texas.

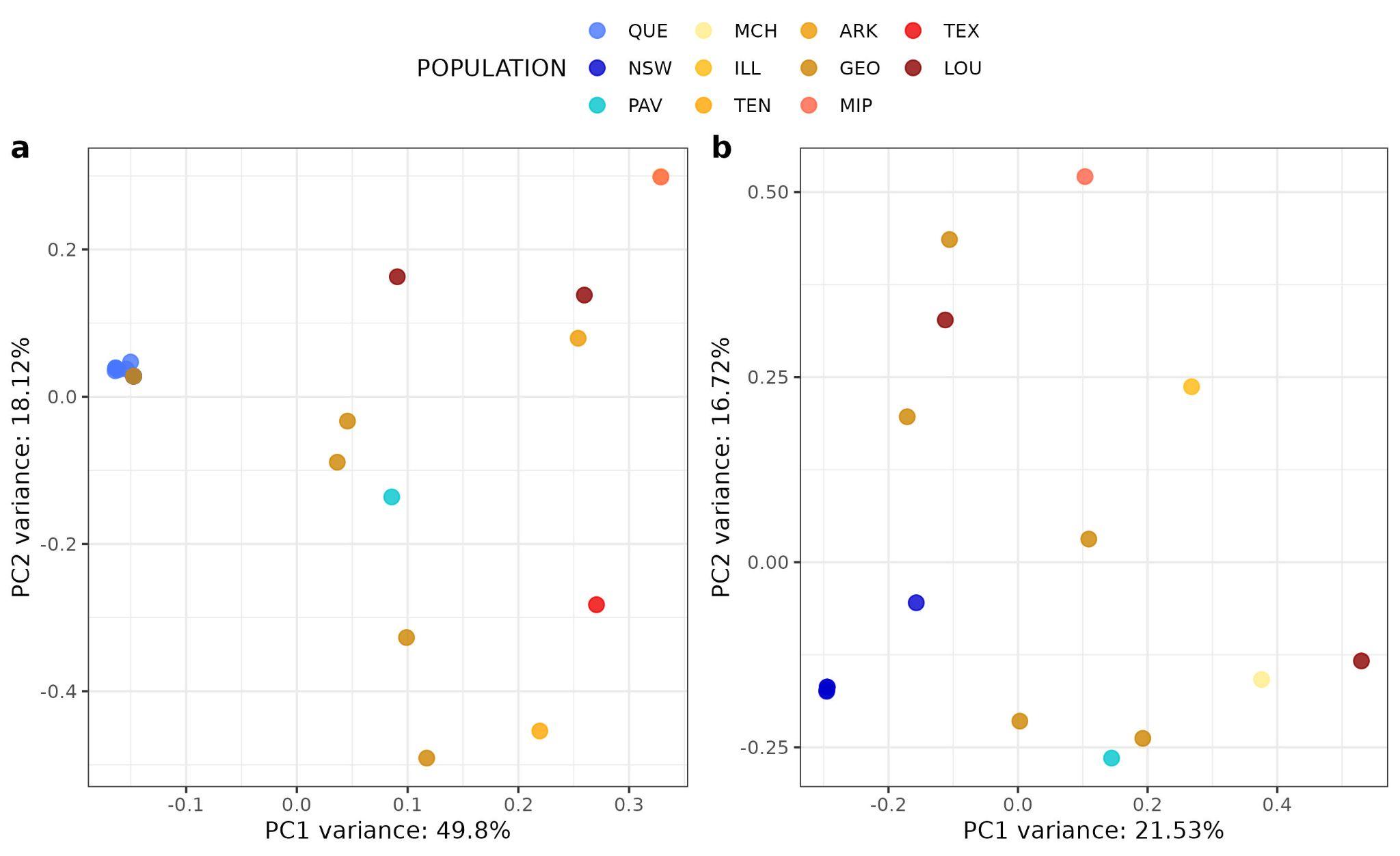

**Fig. S5. PCA on mitochondrial (a) and Wolbachia (b) variants.** Queensland, QUE; New South Wales, NSW; SIC, Sichuan; Pavia, PAV; Michigan, MCH; Illinois, ILL; Tennessee, TEN; Arkansas, ARK; Mississippi, MIP; Texas, TEX; Georgia, GEO; Louisiana, LOU. In Fig. S2a, several samples from Australia are placed in the same position since the variants present in those samples were identical.

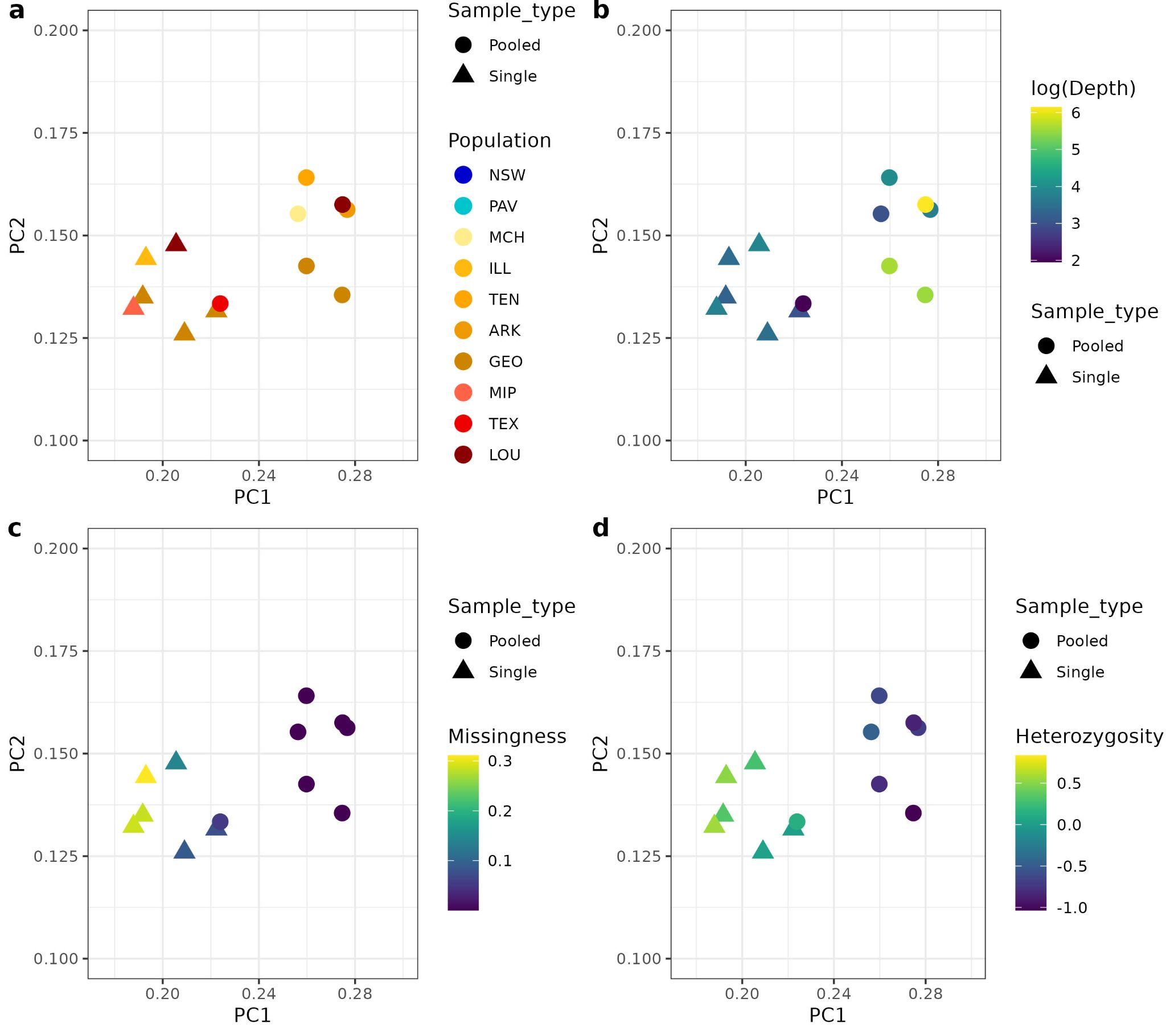

**Fig. S6. PCA of nuclear variants from USA samples.** We performed exploratory analyses using sample metadata to understand the genetic distribution of the US samples in the PCA, including (**a**) geographical information and sample type; (**b**) per-sample depth and sample type; (**c**) per-sample missingness and sample type; and (**d**) heterozygosity estimated by inbreeding coefficient and sample type. Queensland, QUE; New South Wales, NSW; Pavia, PAV; Michigan, MCH; Illinois, ILL; Tennessee, TEN; Arkansas, ARK; Mississippi, MIP; Texas, TEX; Georgia, GEO; Louisiana, LOU.

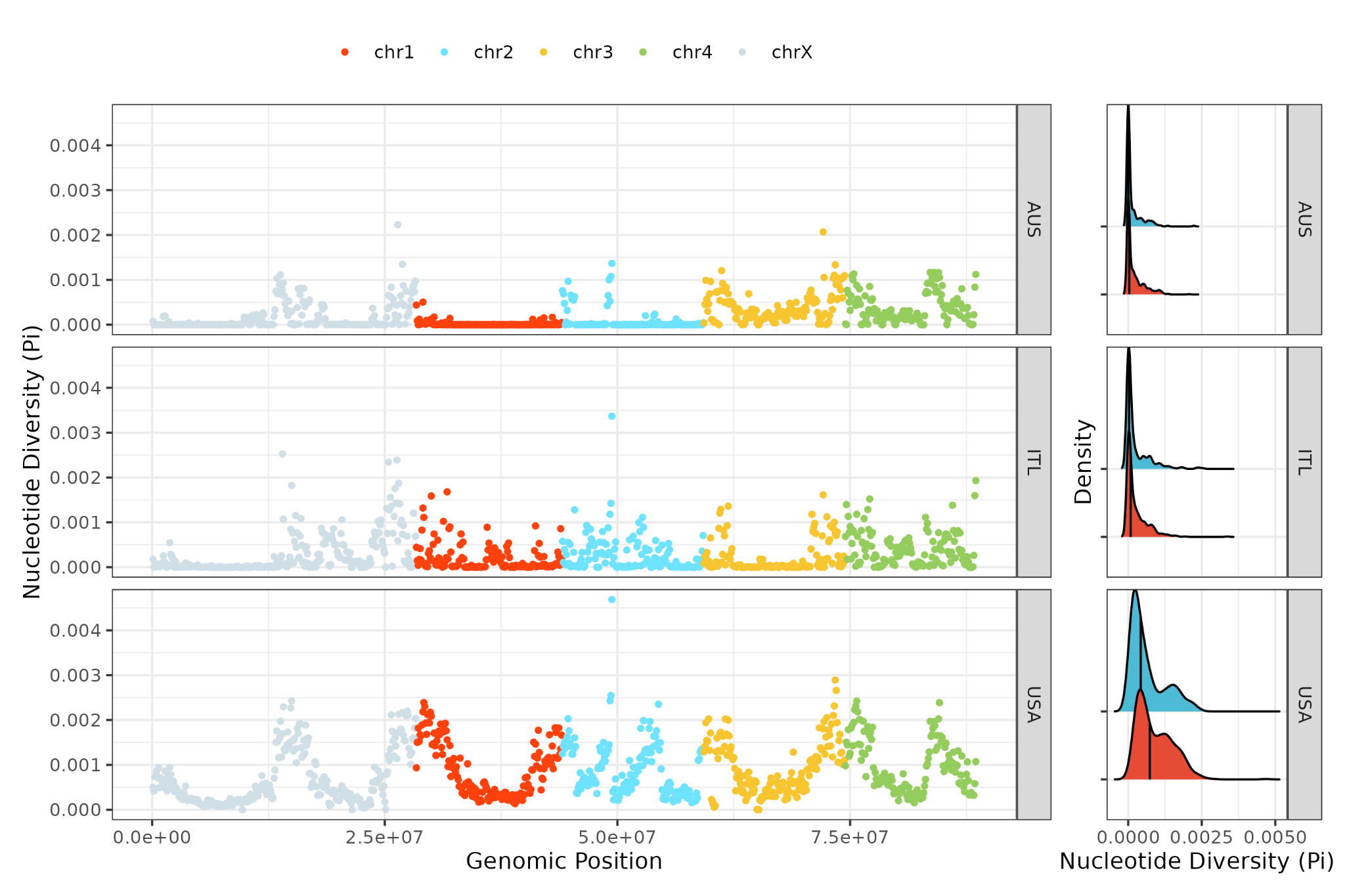

**Fig. S7. Genome-wide patterns of nucleotide diversity (Pi) between USA, Italian and Australian samples.** The colour scale corresponds to each chromosome. Density plots show the distribution of Pi values for the sex-linked (blue) and autosomal (red) scaffolds. The median Pi value for each type of chromosome is shown (solid vertical line).

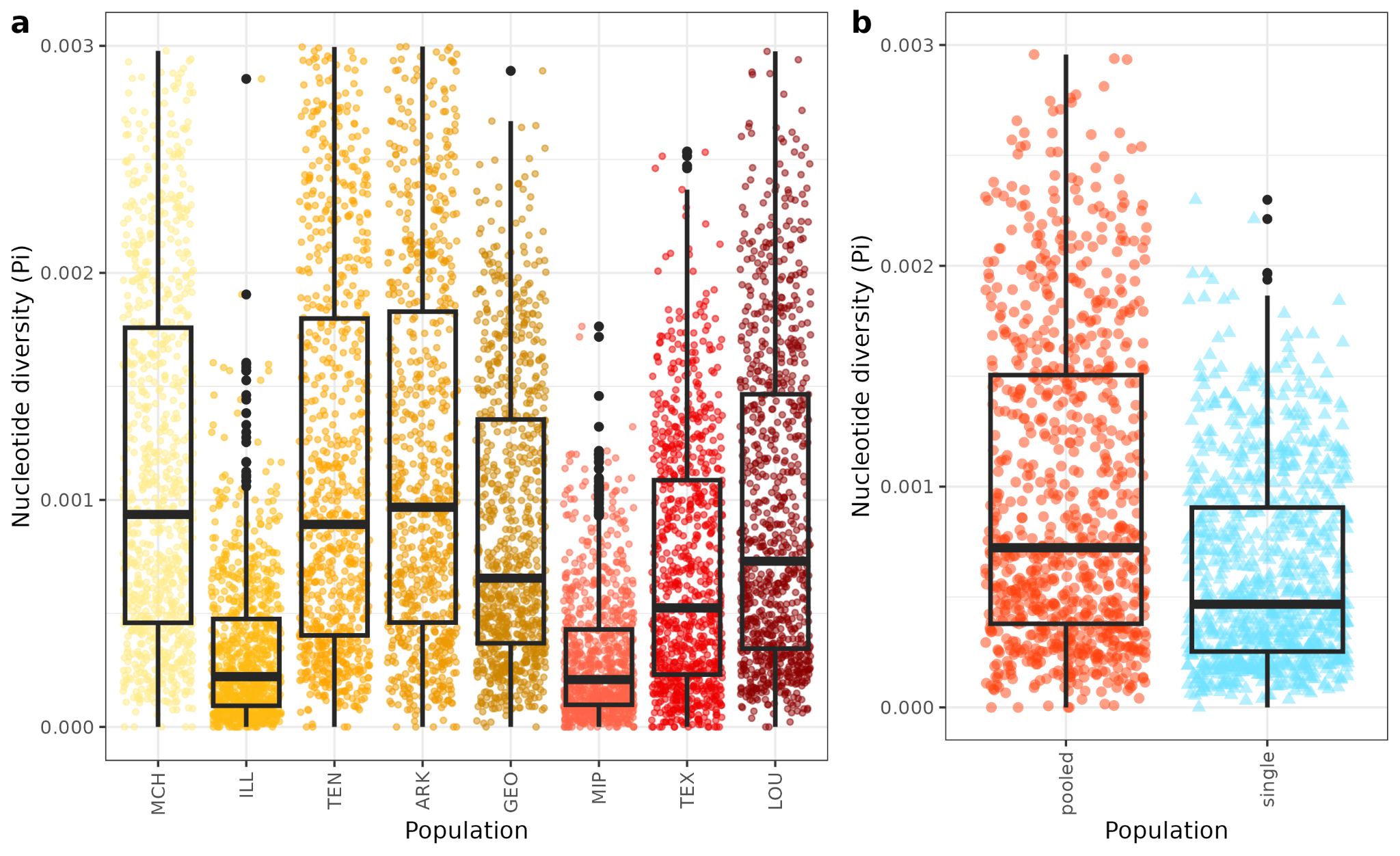

**Fig. S8. Nucleotide diversity within the USA.** (**a**) Boxplots of the nucleotide diversity (Pi) per USA population, which are ranked from northern states (left) to southern states (right). (**b**) Boxplot of the Pi within the USA samples divided by sample type: pooled microfilaria and single worms. These data suggest that some populations are more diverse than others in the USA, however, this interpretation is somewhat confounded by the fact that there are different sample types per population, and that pooled samples showed higher diversity than single samples. Michigan, MCH; Illinois, ILL; Tennessee, TEN; Arkansas, ARK; Mississippi, MIP; Texas, TEX; Georgia, GEO; Louisiana, LOU.
